# Disease heritability enrichment of regulatory elements is concentrated in elements with ancient sequence age and conserved function across species

**DOI:** 10.1101/420166

**Authors:** Margaux L.A. Hujoel, Steven Gazal, Farhad Hormozdiari, Bryce van de Geijn, Alkes L. Price

## Abstract

Regulatory elements, e.g. enhancers and promoters, have been widely reported to be enriched for disease and complex trait heritability. We investigated how this enrichment varies with the age of the underlying genome sequence, the conservation of regulatory function across species, and the target gene of the regulatory element. We estimated heritability enrichment by applying stratified LD score regression to summary statistics from 41 independent diseases and complex traits (average *N* =320K) and meta-analyzing results across traits. Enrichment of human enhancers and promoters was larger in elements with older sequence age, assessed via alignment with other species irrespective of conserved functionality: enhancer elements with ancient sequence age (older than the split between marsupial and placental mammals) were 8.8x enriched (vs. 2.5x for all enhancers; p = 3e-14), and promoter elements with ancient sequence age were 13.5x enriched (vs. 5.1x for all promoters; p = 5e-16). Enrichment of human enhancers and promoters was also larger in elements whose regulatory function was conserved across species, e.g. human enhancers that were enhancers in ≥5 of 9 other mammals were 4.6x enriched (p = 5e-12 vs. all enhancers). Enrichment of human promoters was larger in promoters of loss-of-function intolerant genes: 12.0x enrichment (p = 8e-15 vs. all promoters). The mean value of several measures of negative selection within these genomic annotations mirrored all of these findings. Notably, the annotations with these excess heritability enrichments were jointly significant conditional on each other and on our baseline-LD model, which includes a broad set of coding, conserved, regulatory and LD-related annotations.

## Introduction

Disease-associated variants and disease heritability have been widely reported to be concentrated in regulatory annotations, such as enhancers and promoters ^1^–^7^. These findings have motivated recent studies of how enhancers and promoters evolve across species ^8^–^11^. Vierstra et al. analyzed DNase I Hypersensitivity Sites (DHS) in humans and mice and reported that human-specific DHS were significantly enriched for disease-and trait-associated variants, despite decreased constraint^8^. Villar et al. analyzed 20 mammalian species and reported that enhancers evolve more rapidly than promoters, and that enhancers were often species-specific whereas promoters were often functionally conserved ^9^. Vermunt et al. and Trizzino et al. analyzed 3-6 primate species and reported that regulatory elements were generally functionally conserved across primates, with higher sequence and function conservation for promoters than for enhancers ^10,11^. However, which enhancers and promoters are most important for disease remains largely unknown. Further investigating which enhancers and promoters are most important for disease would improve our biological understanding of disease architectures.

Here, we characterize the contribution of enhancers and promoters to disease heritability based on sequence age, conserved function across species, and gene function of the target gene. We achieve this by constructing new annotations using enhancers and promoters previously identified in liver tissue using 10 high-quality genomes (humans and 9 other mammalian species ^9^) and applying stratified LD score regression with the baseline-LD model ^6,7^ to summary association statistics from 41 independent diseases and complex traits (average *N* =320K). We find that disease heritability enrichment is concentrated in enhancers and promoters with ancient sequence age and conserved function across species, as well as promoters of loss-of-function intolerant genes from the Exome Aggregation Consortium (ExAC)^12^. The mean value of several measures of negative selection within these genomic annotations mirrored all of these findings, with larger heritability enrichments for annotations under stronger negative selection.

## Methods

### Enhancer and promoter annotations

Our goal is to understand the role of human enhancers and promoters in the genetic architecture of diseases and complex traits. We first annotated regions as enhancers and promoters using previously identified enhancer and promoter regions that were enriched for histone marks (H3K27ac and H3K4me3) in at least 2 of 4 human liver tissue samples ^9^. We merged any overlapping annotations, resulting in enhancer and promoter regions with mean segment lengths of 3.4kb and 4.3kb, respectively. In total, 3.3% of common variants lie within enhancers and 1.5% within promoters (Table 1).

**Table 1:** **Annotations analyzed in main analyses.** We report the proportion of common SNPs (MAF 0.05) and mean segment length in base pairs (BP) for each annotation. For the count annotations (enhancer conservation count and promoter conservation count), we report here the corresponding binary annotations, conserved enhancer and conserved promoter. Mean segment length is computed after merging overlapping elements.

| Annotation | Prop. SNPs | Prop. enhancer/promoter | Mean segment length (BP) |
| --- | --- | --- | --- |
| Enhancer | 0.0332 |  | 3,362 |
| Promoter | 0.0152 |  | 4,308 |
| Ancient enhancer | 0.0052 | 0.1557 | 191 |
| Ancient promoter | 0.0042 | 0.2765 | 159 |
| Conserved enhancer | 0.0055 | 0.1649 | 4,962 |
| Conserved promoter | 0.0080 | 0.5251 | 4,502 |
| Promoter of ExAC gene | 0.0025 | 0.1640 | 4,549 |

### Sequence age annotations

We constructed genomic annotations based on genome-wide alignments of 100 vertebrates ^13^. Each region of the human genome has an associated score between 1 and 19 based on the number of key ancestral nodes in the tree of vertebrates that it aligned to (1st root=human; 19th root=vertebrates); younger regions were assigned smaller scores, whereas older regions were assigned larger scores. Most regions were assigned a precise age (one score), but some regions were assigned an age interval (range of scores) or an inconsistent age. Regions with inconsistent age were removed.

We assigned an age to each SNP lying in an enhancer or promoter based on the corresponding region of the genome (start location ≤ SNP location < end location). We removed SNPs in regions in which the alignment at the 19th root was uncertain and assigned the maximum inferred age for SNPs in regions with an age interval. We categorized the ages as post-eutheria split (1-11; young), eutheria (12; intermediate), and pre-eutheria split (13-19; ancient), with approximately one third of enhancers and promoters falling in each of these age bins. Pre-eutheria split (or ancient sequence age) means the sequence has an age older than the split between placental and marsupial mammals (>160 million years old ^14,15^).

We also analyzed enhancers and promoters from the baseline model ^6^: Hoffman enhancers ^16^ and UCSC promoters ^17^ that are not tissue-specific, unlike Villar et al. enhancers and promoters (Table S2). We annotated the Hoffman enhancers and UCSC promoters as post-eutheria split (1-11; young), eutheria (12; intermediate), and pre-eutheria split (13-19; ancient) using the same procedure as for the Villar et al. enhancer and promoter annotations.

### Conserved function annotations

We annotated human enhancers and promoters according to their conserved function, assessed via how many of 9 other mammalian species (with high quality genomes) assayed by ref. ^9^ had shared regulatory functionality. Human-specific and highly conserved enhancers and promoters were defined as elements with conserved function in 0 or 9 of the 9 mammals, respectively. We denote conserved enhancers and promoters as elements with conserved function in at least 5 of the 9 mammals. We constructed 6 categorical annotations (3 for promoters and 3 for enhancers): each enhancer and promoter was annotated with the conservation count (CC) in other species (CC= 0, 1,…, 9; both align and have functional conservation), the mapped count in other species (0, 1,…, 9; align but no functional conservation) and the missing count in other species (0, 1,…, 9). We introduced 20 binary annotations (10 for promoters and 10 for enhancers) reflecting the 10 possible values of CC (0, 1,…, 9).

We computed the conservation count, mapped count and missing count of all elements (prior to merging) and then merged information across overlapping elements. For elements that overlapped, for each of conserved, mapped, and missing count, we computed the union of each count across species; this implies that these small proportions of the genome where two or more elements overlap could get conservation, mapped, and missing counts that add up to more than 9.

### Gene function annotations

To assess how the target gene may impact the role of a promoter in disease architecture, we annotated promoters based on whether they were a promoter of an ancient gene (P1-P10 genes ^18^–^20^ which emerged before the vertebrates split ≈500 million years ago ^21^), a loss-of-function intolerant gene from ExAC (ExAC gene ^12^; 3,230 such genes; Table 1), or a gene with a mouse ortholog (identified in hg38; we assume that gene names remain consistent across builds) ^22^.

We obtained the coordinates of all TSSs and associated genes ^23^. We calculated the mid-point of each merged promoter and determined whether the closest transcription start site (TSS) within 5 kb to the midpoint corresponded to a gene in the specified gene set.

### Heritability enrichment and standardized effect size (*τ*^*\**^) metrics

In order to estimate the heritability enrichment of an annotation, we ran stratified LD score regression (S-LDSC)^6,7^ using 1000 Genomes as the LD reference panel ^24^. Consider *C* binary or continuous-valued annotations (*a*_1_,…, *a*_*C*_), denote *a*_*c*_(j) the annotation value of SNP j for annotation *c*, and assume that the variance of per normalized genotype effect sizes linearly depends on the *C* annotations: Var(*β*_*j*_)=∑_*c*_ *a*_*c*_(j)τ_*c*_, where *τ*_*c*_ is the per-SNP contribution of one unit of the annotation c to heritability (jointly modeled with all other annotations). S-LDSC estimates *τ*_*c*_ using the summary statistic for a SNP 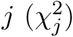 via the following equation:

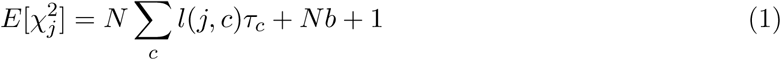

where *N* is the sample size of the GWAS, *b* quantifies confounding biases ^25^, and 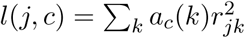 is the LD score of SNP *j* to annotation *c* where r_*jk*_ is the correlation between SNPs *j* and *k*. S-LDSC estimates two metrics quantifying the role of a functional region in diseases and complex traits. First, it estimates the heritability enrichment of binary annotations, defined as the proportion of heritability explained by SNPs in the annotation divided by the proportion of SNPs in the annotation. The enrichment of annotation *c* is estimated as

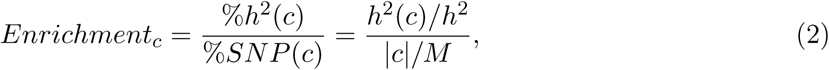

where *h*^2^(c) is the heritability causally explained by common SNPs in annotation *c, h*^2^ is the heritability causally explained by common SNPs, |*c*| is the number of common SNPs that lie in the annotation, and M is the number common SNPs (in our analyses M=5,961,159 SNPs, see below). A value greater than one would indicate a functional annotation is enriched for trait heritability or the proportion of heritability explained is greater than one would expect given the size of the annotation.

Standardized effect size 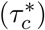 was previously defined ^7^ as the proportionate change in per-SNP heritability associated with a one standard deviation increase in the value of the annotation, conditional on the other annotations in the model; 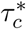 quantifies effects that are unique to the focal annotation, unlike heritability enrichment^6,7,26^. In detail,

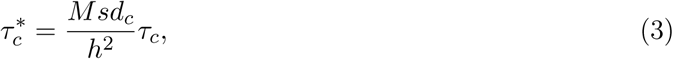

where *sd*_*c*_ is the standard deviation of annotation *c*.

Regression SNPs (the SNPs used by S-LDSC to estimate *τ*_*c*_ from marginal association statistics) were obtained from the HapMap Project phase 3; these SNPs are considered to be well-imputed SNPs. SNPs with marginal association statistics larger than 80 or 0.001N and SNPs that are in the major histocompatibility complex (MHC) region were excluded from all analyses. Reference SNPs (the SNPs used by S-LDSC to compute LD scores) were defined as the set of 9,997,231 biallelic SNPs with minor allele count greater or equal than five in the set of 489 unrelated and outbred European samples ^27^ from phase 3 of 1000 Genomes Project (1000G) ^24^. We note that regression SNPs tag potentially causal reference SNPs via LD scores computed using reference SNPs^6,7^. Heritability SNPs (the SNPs used by S-LDSC to compute *h*^2^, *h*^2^(*c*), | *c* |and *sd*_*c*_) were defined as the 5,961,159 common variants (MAF ≥ 0.05) in the set of reference SNPs. Using the LD score for each annotation and the marginal statistics obtained from the trait phenotypes, we computed the heritability enrichment and 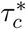 for each annotation.

In all analyses we included the enhancer and promoter annotations ^9^ as well as a broad set of 75 functional annotations from the baseline-LD (v1.1) model ^7,26^, which include functional annotations (i.e. coding, intron, DHS, …), 10 MAF bins and 6 LD-related annotations (Table S2). We note that the inclusion of MAF-and LD-related annotations implies that the expected causal heritability of a SNP is a function of MAF and LD. We meta-analyzed results across a previously described set ^26^ of 41 independent diseases and complex traits (average *N* =320K, computed using largest data set for each trait); for six traits we analyzed two data sets (genetic correlation > 0.9), leading to a total of 47 data sets analyzed (Table S3). We performed random-effects meta-analyses across traits using the R package rmeta.

Reported enrichment estimates are based on a random-effects meta-analysis of enrichment estimates for each trait. The p-value for enrichment is computed by meta-analyzing 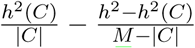 across the traits using a random-effects model (as enrichment is not normally distributed ^6^) and testing the null hypothesis that this difference is 0. For enhancer conservation count and promoter conservation count we calculate enrichment for bins of this categorical annotation ^7^.

For each new annotation, we ran S-LDSC conditional on the enhancer and promoter annotations as well as the baseline-LD model. For each annotation type (sequence age, conserved function, and gene function) we derived a joint model by running S-LDSC with the full set of annotations of that type conditional on the enhancer and promoter annotations and the baseline-LD model. (For the sequence age analysis this set consisted of 4 annotations: young enhancer/promoter and ancient enhancer/promoter; we removed the intermediate enhancer/promoter to avoid linear dependence between the annotations. For the conserved function model this set consisted of 8 annotations: human-specific enhancer/promoter, highly conserved enhancer/promoter, enhancer/promoter CC and enhancer/promoter missing count; we removed enhancer/promoter mapped count to avoid linear dependence between the annotations. For the gene function model this set consisted of 3 annotations: promoter of an ancient gene, promoter of ExAC gene, and promoter of a gene with a mouse ortholog.) We then iteratively removed the least statistically significant annotation (excluding annotations in the baseline-LD model as well as the enhancer and promoter annotation) until each remaining annotation was significant (after correction for multiple testing) ^7^. To produce a combined joint model, we combined the significant annotations sequence age, conserved function, and gene function annotations into a single model and again iteratively removed the least statistically significant annotation until each annotation remained significant (after correction for multiple testing).

In order to determine whether a subset of enhancers and promoters were particularly enriched as compared to all enhancers or promoters, we computed the enrichment difference between an annotation A and a subset a. For each trait, we computed the difference in enrichment between the annotations (Δ) and the standard error for this difference (using block-jackknife) and then meta-analyzed results across 41 traits using random-effects meta-analysis. In order to compute a p-value for the difference in enrichment we computed the normally distributed quantity 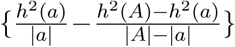 as well as its standard error for each trait (using block-jackknife), and then meta-analyzed results across traits. We then tested the null hypothesis that this difference is 0; this test assesses whether the per-SNP heritability within annotation A is different within a than outside a. This test is a natural extension of the approach used to assess statistical significance of enrichment^6^.

We computed the proportion of enrichment for an annotation A, attributable to a subset a. The proportion of enrichment for A attributable to a is defined as

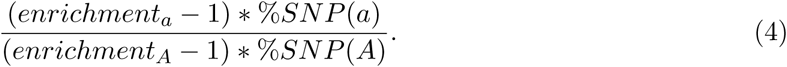

If *enrichment*_*a*_ = *enrichment*_*A*_, then the proportion of enrichment for A attributable to a is just the proportion of A in subset a. We computed this quantity for each trait, used block-jacknife to compute standard errors, and meta-analyzed results across 41 traits.

### Negative selection metrics

We quantified the strength of negative selection within these annotations by computing the mean value of several measures of negative selection and computing the standard error using block-jackknife with 200 equally sized blocks of adjacent SNPs within the annotations (all measures are annotations in the baseline-LD model; Table S2). First, we computed the proportion of common SNPs with GERP++ rejected substitutions (RS) score 4 (GERP RS 4, binary annotation) within the baseline-LD annotations ^7,28^. This score is equal to the difference between the neutral and observed substitution rates and reflects the intensity of constraint at a given genomic location, such that a larger score is indicative of stronger negative selection. Second, we computed the mean background selection statistic (BSS = 1-McVicker B statistic ^29^) (at common SNPs); a BSS value close to 1 indicates that background selection resulted in near complete removal of diversity whereas a value close to 0 indicates little effect ^7^. Third, we computed the proportion of common SNPs conserved across mammals ^6,30^; regions that are conserved across mammals are likely to be critical, as mutations were not tolerated. Fourth, we computed the mean MAF-adjusted predicted allele age (at common SNPs); on average, recent variants are more deleterious ^7,31^. Fifth, we computed mean nucleotide diversity ^32^ (at common SNPs); variants that lie in regions with low nucleotide diversity are more likely to be deleterious ^7,33^.

## Results

### Disease enrichment is concentrated in regulatory elements with ancient sequence age

We focused our analyses on enhancer and promoter elements that were previously annotated based on H3K27ac and H3K4me3 marks assayed in human liver ^9^ (Table 1). To assess the disease enrichment of these elements, we applied S-LDSC with the baseline-LD model ^6,7^ to summary statistics from 41 independent diseases and complex traits (average *N* =320K; Table S3) and meta-analyzed results across traits. We observed significant heritability enrichment for both enhancers (2.6x, p= 3e-12) and promoters (4.6x, p= 3e-17) (Table S4A), consistent with previous studies of disease enrichment of regulatory elements ^1^–^7^. Based on significance of regression coefficients we determined that the promoter annotation (but not the enhancer annotation) provides unique information conditioned on the baseline-LD model (p= 0.007; Table S4A), which includes a broad set of regulatory annotations (Table S2). Analyses of highly reproducible enhancer and promoter annotations (reproduced in all four tissue samples from ref. ^9^) produced similar results (Table S4B).

We annotated enhancer and promoter regions according to their underlying sequence age, assessed via genome-wide alignment of 100 vertebrates irrespective of conserved functionality ^13^. Each region of the human genome had an associated score between 1 and 19 based on the number of key ancestral nodes in the tree of vertebrates that it aligned to. We classified enhancer and promoter regions as having a young (1-11), intermediate (12), or ancient (13-19) sequence age (see Methods); different regions within the same enhancer or promoter may be assigned different sequence ages. Ancient sequence age means the sequence is older than the split between marsupial and placental mammals (>160 million years old ^14,15^); 16% of enhancer SNPs were annotated as ancient enhancer, and 28% of promoter SNPs were annotated as ancient promoter (Table 1). The ancient enhancer and ancient promoter annotations were only weakly correlated with annotations from the baseline-LD model (Figure S1 and Table S1).

To assess how the disease enrichment of enhancers and promoters varies with sequence age, we repeated our S-LDSC analysis with each of the 6 age-specific annotations (young, intermediate, or ancient; enhancer or promoter) included in turn, in addition to baseline-LD + enhancer + promoter annotations. We observed the strongest enrichments for ancient enhancers and ancient promoters (Table S5). We constructed a joint sequence age model by retaining only the age-specific annotations that remained significant (after correction for multiple testing) when conditioned on the baseline-LD + enhancer + promoter annotations ^7^; only the ancient enhancer and ancient promoter annotations were jointly significant. Ancient enhancers were 9.3x enriched, compared to 2.7x for all enhancers (p=4e-15 for difference), and ancient promoters were 14.3x enriched, compared to 4.9x for all promoters (p=2e-18 for difference) (Figure 1A and Table S6A). We note that enrichment estimates can change slightly depending on the set of annotations included in the model ^6,7^ (see Combined joint model section for enrichment estimates reported in Abstract). Although ancient enhancers comprise only 16% of enhancers, they contribute 59% (s.e. 5%) of all enhancer enrichment. Analogously, although ancient promoters comprise only 28% of promoters, they contribute 82% (s.e. 4%) of all promoter enrichment.

**Figure 1:**
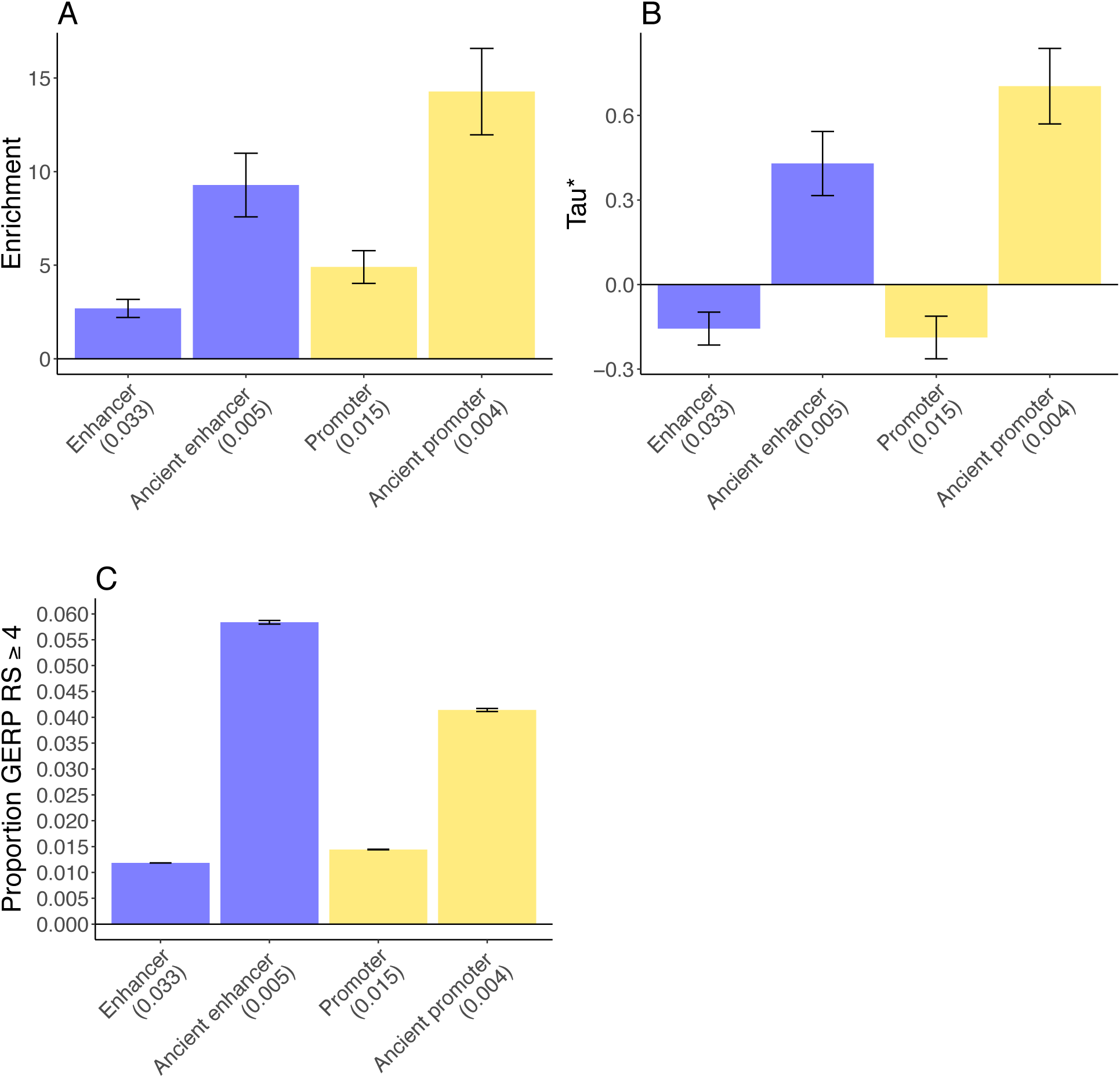
Disease enrichment of ancient enhancers and ancient promoters in sequence age model. We report results for sequence age annotations that are jointly significant conditional on the baseline-LD model and enhancer and promoter annotations (Bon-ferroni p = 0.05/4 = 0.0125). (A) Heritability enrichment and (B) *τ*^*^ estimates (± 1.96 standard error); results are meta-analyzed across 41 traits. (C) Proportion of common SNPs within annotations with GERP RS ≥ 4^7,28^ (± 1.96 standard error). We report the proportion of common SNPs (MAF ≥ 0.05) for each annotation. Numerical results are reported in Table S6.

Both ancient enhancers and ancient promoters were uniquely informative for disease heritability conditional on the baseline-LD + enhancer + promoter annotations, as quantified by *τ*^*^ (the proportionate change in per-SNP heritability associated with an increase in the value of the annotation by one standard deviation, conditional on other annotations included in the model ^7^) (Figure 1B and Table S6B). Specifically, we estimated large and highly significant values of *τ*^*^ for both ancient enhancers (*τ*^*^ = 0.43, p=1e-13) and ancient promoters (*τ*^*^ = 0.70, p=9e-25). In particular, these *τ*^*^ values were larger than the analogous *τ*^*^ values that we recently estimated for both LD-related annotations ^7^ and molecular QTL annotations ^26^, implying a substantial improvement in our understanding of which regulatory elements contribute to disease heritability. The slightly but significantly negative value of *τ*^*^ for (all) enhancers and (all) promoters indicates conditional depletion for enhancers and promoters that do not have ancient sequence age (Figure 1B and Table S6B).

We quantified the mean strength of negative selection within each of the annotations from Figure 1A. We first calculated the proportion of common SNPs with GERP++ rejected substitutions (RS) score ≥4 (GERP RS≥ 4) ^7,28^. The GERP RS score reflects the difference between the neutral and observed substitution rates and thus reflects the intensity of constraint at a given genomic location, such that a larger score is indicative of stronger negative selection. We determined that the stronger disease enrichment for ancient enhancers and ancient promoters is mirrored by the larger proportion of variants in these annotations with GERP RS 4, reflecting stronger negative selection (Figure 1C and Table S6C). We observed similar patterns for 4 other measures of negative selection ^7^: a background selection statistic (BSS) equal to 1-McVicker B statistic ^29^; sequence conservation across 29 mammals ^30^; predicted allele age ^7^; and nucleotide diversity ^32^ (Table S6C). However, as noted above, ancient enhancers and ancient promoters were uniquely informative for disease heritability conditional on the baseline-LD model, which includes all of these measures of negative selection.

We performed three secondary analyses to assess the robustness of our results. First, we repeated the analysis of Figure 1A-B by adding a binary annotation defined by ancient sequence age (irrespective to enhancer or promoter status) to the model; this annotation was not conditionally informative for disease heritability as quantified by *τ*^*^, and its addition to the model did not significantly change our results (Table S7). Second, we repeated the analysis of Figure 1A-B by using the enhancer and promoter annotations from the baseline-LD model ^7^ (instead of ref.^9^; Table S2). We obtained similar results, with much stronger enrichments for ancient enhancers and ancient promoters (Table S8). (We used the enhancer and promoter annotations from ref. ^9^ in our main analyses so that we could integrate annotations based on ancient sequence age and conserved function into a combined joint model; see below.) Third, we repeated the analysis of Figure 1A-B by including 500bp flanking regions around each of the annotations from Figure 1A-B, to guard against bias due to model misspecification ^6^. We confirmed that this did not significantly change our results (Table S9).

### Disease enrichment is concentrated in regulatory elements with conserved function

We annotated human enhancers and promoters according to their conserved function, assessed via how many of 9 other mammalian species assayed by ref. ^9^ had shared regulatory functionality. Each enhancer and promoter was annotated with the conservation count (CC) in other species (CC= 0, 1,…, 9). We constructed both integer-valued “conservation count” (value of CC) and binary “conserved” (CC 5) annotations (see Methods and Table 1). A large proportion of annotated enhancers were functionally human-specific (40% human-specific (CC= 0) vs. 2% highly conserved (CC= 9)), whereas promoters were more functionally conserved (19% human-specific vs. 15% highly conserved) (Table S12). Accordingly, 53% of promoters were conserved promoters, whereas only 16% of enhancers were conserved enhancers (Table 1). The enhancer conservation count and promoter conservation count annotations were only weakly correlated with annotations from the baseline-LD model, but moderately correlated with the ancient enhancer and ancient promoter annotations (Figure S1 and Table S1).

To assess how the disease enrichment of enhancers and promoters varies with conserved function, we performed S-LDSC analyses with each of 10 conserved-function-specific annotations (conservation count, highly conserved, human-specific, mapped count, missing count (see Methods); enhancer or promoter) included in turn, in addition to baseline-LD + enhancer + promoter annotations. We observed the strongest enrichments for highly conserved enhancers and highly conserved promoters, and also observed that while human-specific promoters were enriched, human-specific enhancers were not (Table S10). We constructed a joint conserved function model by retaining only the conserved-function-specific annotations that remained significant (after correction for multiple testing) when conditioned on the baseline-LD + enhancer + promoter annotations ^7^; only the enhancer conservation count and promoter conservation count annotations were jointly significant. Because enrichment is not defined for annotations with value 0-9, we estimated the enrichment of the corresponding binary annotations (conserved enhancer and conserved promoter) in the joint model. Conserved enhancers were 4.6x enriched, compared to 2.4x for all enhancers (p=3e-12 for difference), and conserved promoters were 5.1x enriched, compared to 4.5x for all promoters (p=0.022 for difference) (Figure 2A and Table S11A). We note that enrichment estimates can change slightly depending on the set of annotations include in the model ^6,7^ (see Combined joint model section for enrichment estimates reported in Abstract). Although conserved enhancers comprise only 16% of enhancers, they contribute 35% (s.e. 2%) of all enhancer enrichment. Analogously, although conserved promoters comprise only 53% of promoters, they contribute 59% (s.e. 2%) of all promoter enrichment.

**Figure 2:**
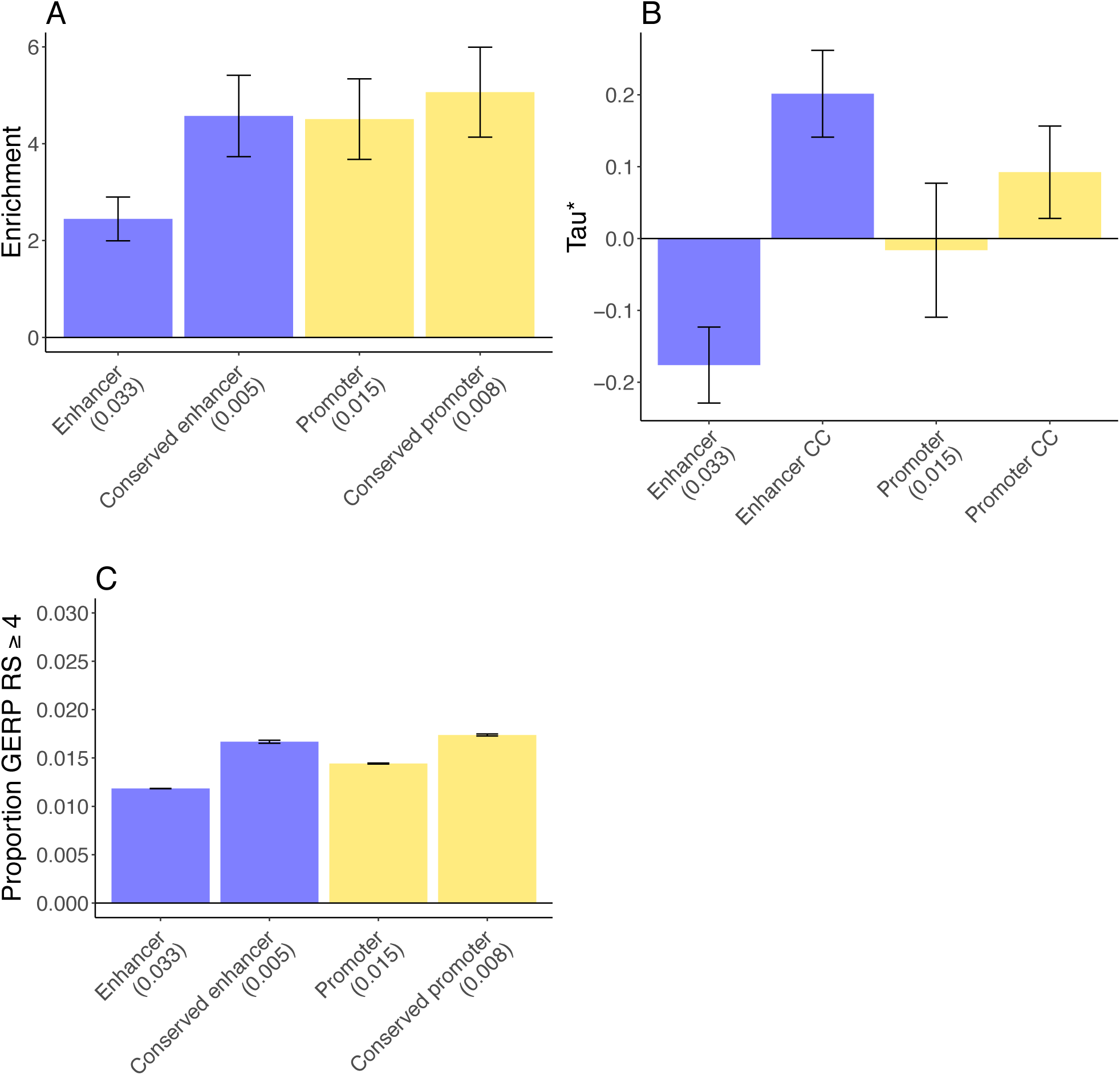
Disease enrichment of conserved enhancers and conserved promoters in conserved function model. We report results for conserved function annotations that are jointly significant conditional on the baseline-LD model and enhancer and promoter annotations (Bonferroni p = 0.05/8 = 0.00625). (A) Heritability enrichment and (B) *τ*^*^ estimates (± 1.96 standard error); results are meta-analyzed across 41 traits. (C) Proportion of common SNPs within annotations with GERP RS ≥ 4^7,28^ (± 1.96 standard error). We report the proportion of common SNPs (MAF ≥ 0.05) for each annotation. Numerical results are reported in Table S11.

Both enhancer conservation count and promoter conservation count were uniquely informative for disease heritability conditional on the baseline-LD + enhancer + promoter annotations, as quantified by *τ*^*^ (Figure 2B and Table S11B). Specifically, we estimated significant values of *τ*^*^ for both enhancer conservation count (*τ*^*^ = 0.20, p=7e-11) and promoter conservation count (*τ*^*^ = 0.10, p=0.005). The significantly negative value of *τ*^*^ for (all) enhancers indicates conditional depletion for enhancers that are not conserved (Figure 2B, Table S11B).

We quantified the mean strength of negative selection within each of the annotations from Figure 2A. We first calculated the proportion of common SNPs with GERP RS 4^7,28^. We determined that the stronger disease enrichments for conserved enhancers and conserved promoters is mirrored by the larger proportion of variants in these annotations with GERP RS 4, reflecting stronger negative selection (Figure 2C and Table S11C). We observed similar patterns for 4 other measures of negative selection (Table S11C). However, as noted above, enhancer conservation count and promoter conservation count were uniquely informative for disease heritability conditional on the baseline-LD model, which includes all of these measures of negative selection.

To further assess how the disease enrichment of enhancers and promoters varies with conserved function, we repeated our S-LDSC analysis with each of 20 binary conservation count annotations (CC= 0, 1,…, 9; enhancer or promoter) jointly included, in addition to baseline-LD + enhancer + promoter annotations. For enhancers, we observed a roughly linear trend whereby enhancers conserved in more mammals are progressively more enriched for heritability (Figure 3A and Table S12A). For promoters, we observed a parabolic trend, similar to the linear trend but with excess heritability for human-specific promoters (Figure 3A and Table S12A).

**Figure 3:**
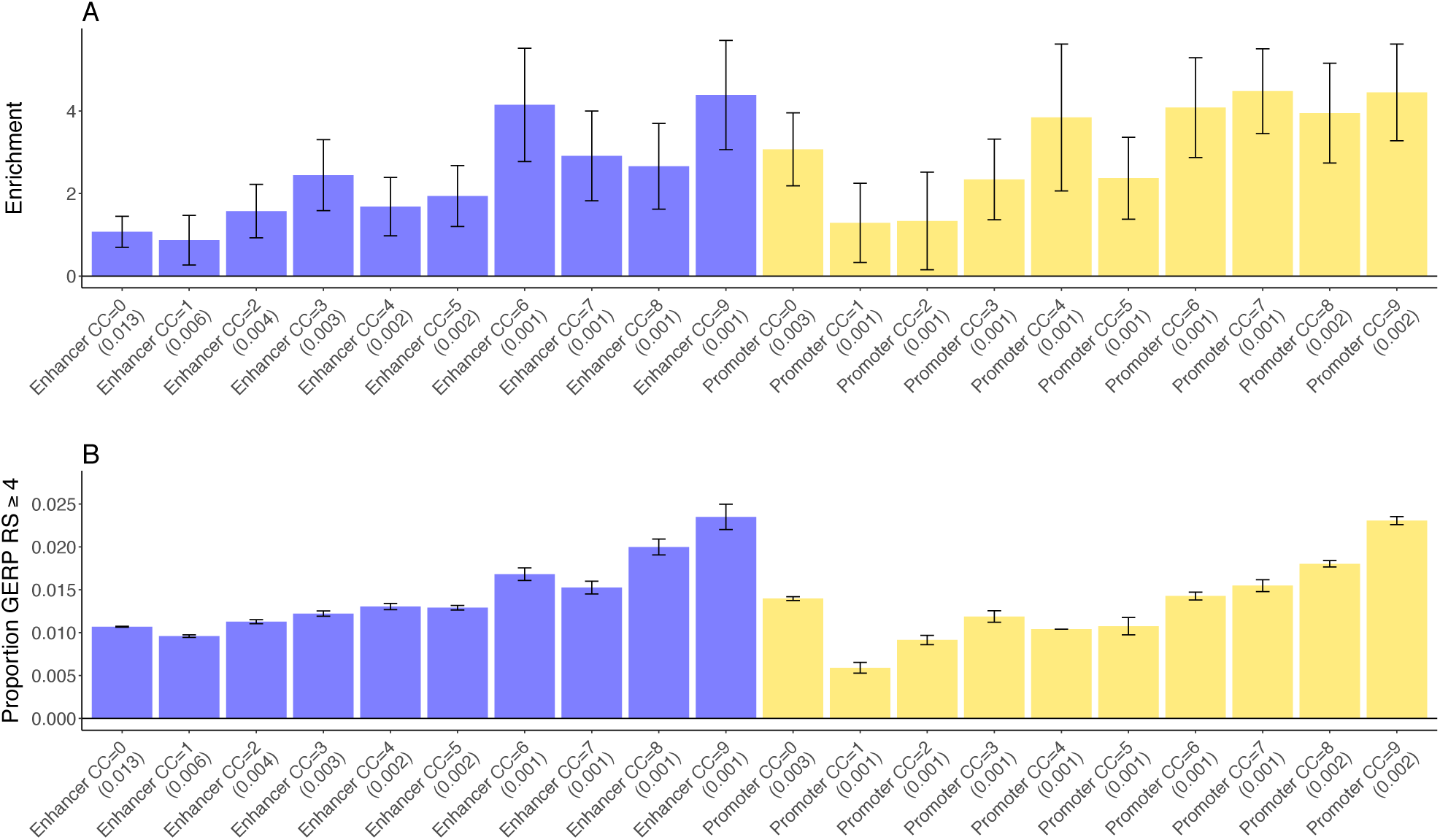
Disease enrichment of enhancers and promoters as a function of conservation count. (A) Heritability enrichment (± 1.96 standard error); results are meta-analyzed across 41 traits. (B) Proportion of common SNPs within annotations with GERP RS ≥4^7,28^ (± 1.96 standard error). We report the proportion of common SNPs (MAF ≥ 0.05) for each annotation. Numerical results are reported in Table S12.

We quantified the mean strength of negative selection within each of the annotations from Figure 3A. We first calculated the proportion of common SNPs with GERP RS 4^7,28^. We determined that the linear disease enrichment trend for enhancers and parabolic disease enrichment trend for promoters (as conservation count increases) is mirrored by the proportion of variants in these annotations with GERP RS 4 (Figure 3B and Table S12B). We observed similar patterns for 4 other S12 measures of negative selection (Table B).

We performed three secondary analyses. First, we repeated the analysis of Figure 2A by replacing the enhancer conservation count and promoter conservation count annotations in the joint model with binary conserved enhancer and conserved promoter annotations, and confirmed that this did not significantly change our results (Table S13). Second, we repeated the analysis of Table S13 by including 500bp flanking regions around each of the annotations from Figure 2A. This did not significantly change our results; the heritability enrichment for conserved enhancer was slightly reduced, but remained highly significant (Table S14). Third, we repeated the analysis of Figure 2B by including human-specific promoters as an additional annotation. While this new annotation was not conditionally significant, the value of *τ*^*^ for the promoter conservation count annotation became larger and more statistically significant (Table S15), consistent with the parabolic trend for promoters in Figure 3A.

### Disease enrichment is concentrated in promoters of loss-of-function intolerant genes

We annotated promoters according to the genes that they regulate (see Methods). In particular, we annotated 16% of promoters as being promoters of the 3,230 ExAC LoF intolerant genes ^12^ (Table 1). The promoter of ExAC gene annotation was only weakly correlated with annotations from the baseline-LD model (Figure S1), but moderately correlated with the ancient promoter and promoter conservation count annotations (Figure S1 and Table S1).

To assess how the disease enrichment of promoters varies with the gene that it regulates, we repeated our S-LDSC analysis with the promoter of ExAC gene annotation included, in addition to baseline-LD + enhancer + promoter annotations. We also analyzed promoter of ancient gene and promoter of gene with mouse ortholog annotations in turn (see Methods). The promoter of ExAC gene annotation produced the strongest enrichment (Table S16), and was the only gene function annotation that remained significant (after correction for multiple testing) in a joint analysis conditioned on the baseline-LD + enhancer + promoter annotations ^7^. Promoters of ExAC genes were 12.4x enriched, compared to 5.1x for all promoters (p=9e-16 for the difference) (Figure 4A and Table S17A). We note that enrichment estimates can change slightly depending on the set of annotations included in the model ^6,7^ (see Combined joint model section for enrichment estimates reported in Abstract). Although promoters of ExAC genes comprise only 16% of promoters, they contribute 39% (s.e. 2%) of all promoter enrichment.

**Figure 4:**
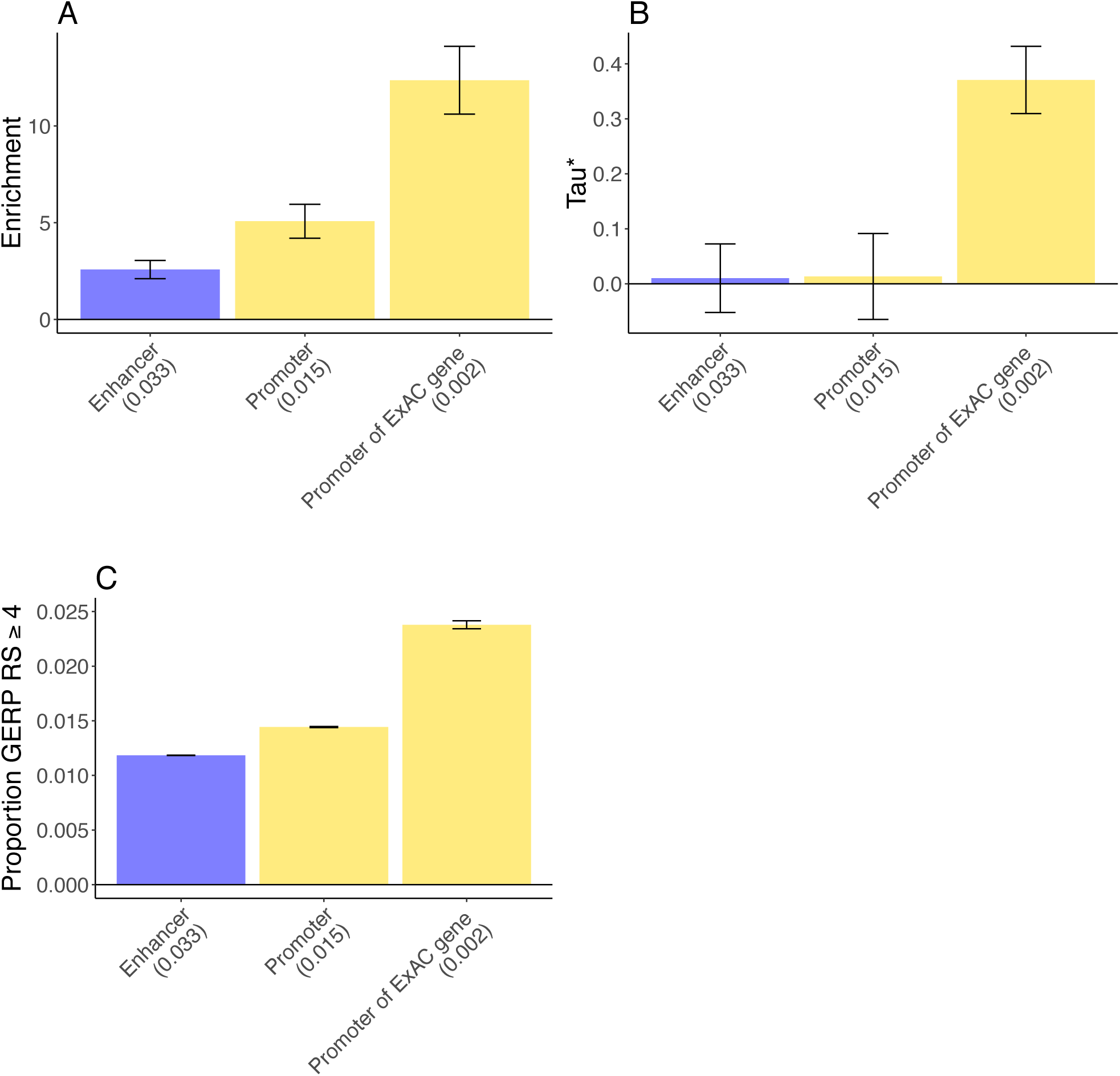
Disease enrichment of promoters of ExAC genes in gene function model. We report results for the gene function annotation that is significant conditional on the baseline-LD model and enhancer and promoter annotations (Bonferroni p = 0.05/3 = 0.0167). (A) Heritability enrichment and (B) *τ*^*^ estimates (±1.96 standard error); results are meta-analyzed across 41 traits. (C) Proportion of common SNPs within annotations with GERP RS ≥4^7,28^ (±1.96 standard error). Numerical results are reported in Table S17. We report the proportion of common SNPs (MAF ≥ 0.05) for each annotation.

Promoters of ExAC LoF intolerant genes were uniquely informative for disease heritability conditional on the baseline-LD + enhancer + promoter annotations, as quantified by *τ*^*^. Specifically, we estimated a large and highly significant value of *τ*^*^ (*τ*^*^ = 0.37, p=2e-32) (Figure 4B and Table S17B).

We quantified the mean strength of negative selection within each of the annotations from Figure 4A-B. We first calculated the proportion of common SNPs with GERP RS ≥ 4^7,28^. We determined that the stronger disease enrichment for promoters of ExAC genes is mirrored by the larger proportion of variants in these annotations with GERP RS ≥ 4, reflecting stronger negative selection (Figure 4C and Table S17C). We observed similar patterns for 4 other measures of negative selection (Table S17C). However, as noted above, promoters of ExAC genes were uniquely informative for disease heritability conditional on the baseline-LD model, which includes all of these measures of negative selection.

We performed two secondary analyses to assess the robustness of our results. First, we repeated the analysis of Figure 4A-B by including 500bp flanking regions around each of the annotations from Figure 4A-B. We confirmed that this did not significantly change our results (Table S18). Second, we repeated the analysis of Figure 4A-B by including two annotations based on fine-mapped expression quantitative trait loci (eQTL): the MaxCPP annotation for all genes and the MaxCPP annotation for ExAC LoF genes only ^26^. Results were little changed, and promoters of ExAC genes were still uniquely informative for disease heritability as quantified by τ ^*\**^ (Table S19).

### Combined joint model

We constructed a combined joint model by including all jointly significant annotations involving sequence age (Figure 1A-B), conserved function (Figure 2B), and gene function (Figure 4A-B) and retaining only the annotations that remained significant (after correction for multiple testing) when conditioned both on each other and on the baseline-LD + promoter + enhancer annotations ^7^. The final joint model included ancient enhancer, ancient promoter, enhancer conservation count, and promoter of ExAC gene annotations. Because enrichment is not defined for annotations with value 0-9, we estimated the enrichment of conserved enhancer in lieu of enhancer conservation count, analogous to above.

Ancient enhancers were 8.8x enriched, compared to 2.5x for all enhancers (p= 3e-14 for difference), and ancient promoters were 13.5x enriched, compared to 5.1x for human promoters (p= 5e-16 for difference) (Figure 5A and Table S20A); these enrichments differed only slightly from the joint sequence age model (Figure 1A). Conserved enhancers were 4.6x enriched (p= 5e-12 for difference vs. all human enhancers); this enrichment differed only very slightly from the joint conserved function model (Figure 2A). Promoters of ExAC genes were 12.0x enriched (p= 8e-15 for difference vs. all promoters); this enrichment differed only very slightly from the joint gene function model (Figure 4A).

**Figure 5:**
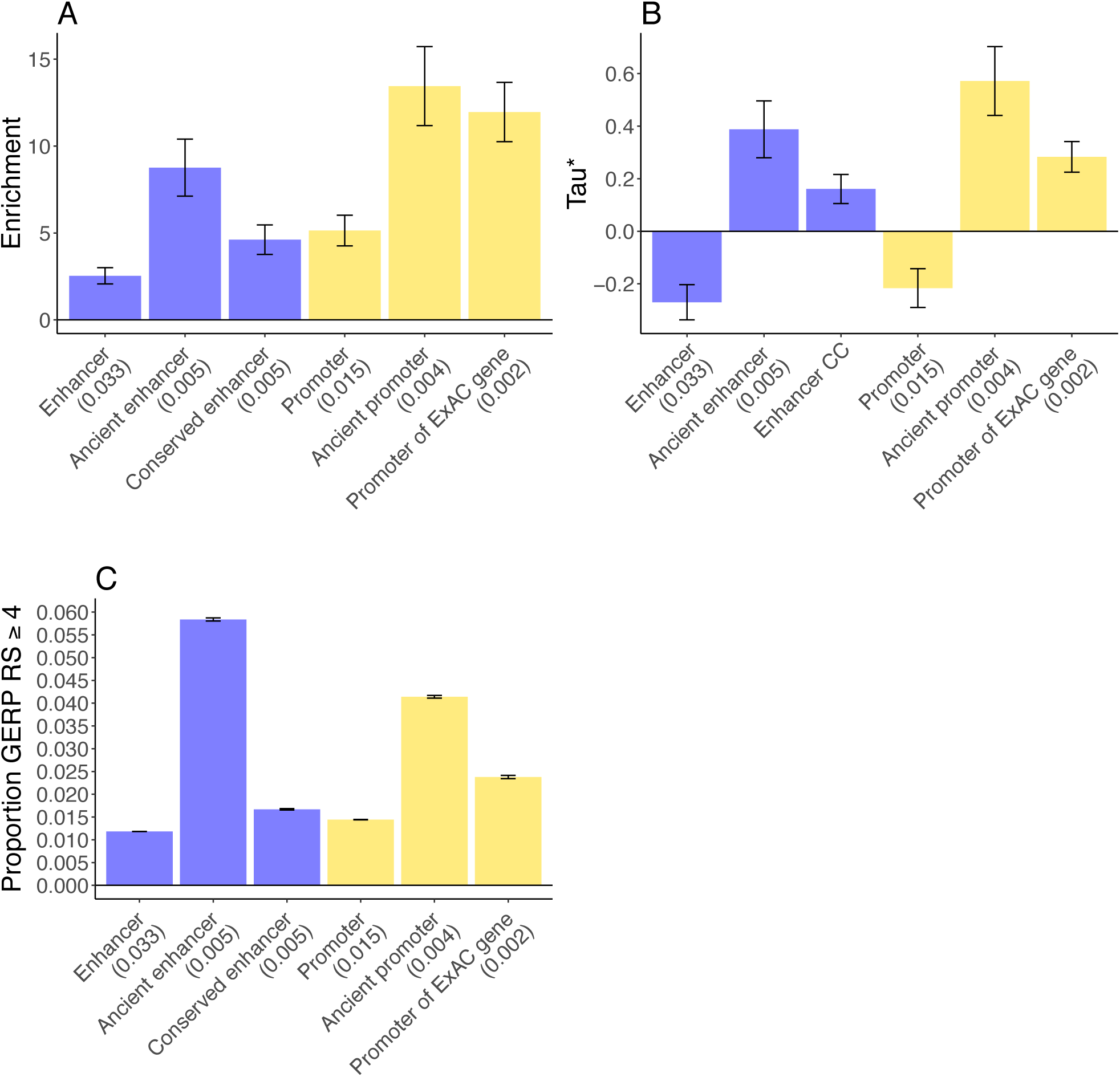
Disease enrichment of annotations in combined joint model. We report results for sequence age, conserved function, and gene function annotations that are jointly significant conditional on the baseline-LD model and enhancer and promoter annotations (Bonferroni p = 0.05/15 = 0.0033). (A) Heritability enrichment and (B) *τ*^*^ estimates (± 1.96 standard error); results are meta-analyzed across 41 traits. (C) Proportion of common SNPs within annotations with GERP RS ≥ 4^7,28^ (± 1.96 standard error). We report the proportion of common SNPs (MAF ≥ 0.05) for each annotation. Numerical results are reported in Table S20.

In the combined joint model, we estimated highly significant values of *τ*^*^ for ancient enhancers (*τ*^*^ = 0.39, p=2e-12), ancient promoters (*τ*^*^ = 0.57, p=1e-17), enhancer conservation count (*τ*^*^ = 0.16, p=1e-8), and promoters of ExAC genes (*τ*^*^ = 0.28, p=2e-21) (Figure 5B and Table S20B). These *τ*^*^ estimates were slightly lower than the corresponding *τ*^*^ estimates from the joint sequence age model (Figure 1B), joint conserved function model (Figure 2B), and joint gene function model (Figure 4B), consistent with correlations between these annotations (Figure S1 and Table S1). Notably, the *τ*^*^ estimates for ancient enhancers and ancient promoters remained larger than the analogous *τ*^*^ values that we recently estimated for LD-related annotations ^7^ and molecular QTL annotations ^26^.

The stronger disease enrichment for ancient enhancers, ancient promoter, conserved enhancers, and promoters of ExAC genes is mirrored by the larger proportion of variants in these annotations with GERP RS ≥ 4 and 4 other measures of negative selection, reflecting stronger negative selection S20 (Figure 5C and Table C), as we previously determined (Figure 1C and Table S6C; Figure 2C S11 and Table C; Figure 4C and Table S17C). However, as noted above, all of these annotations were uniquely informative for disease heritability conditional on the baseline-LD model, which includes all of these measures of negative selection.

We performed three secondary analyses to assess the robustness of our results. First, we repeated the analysis of Figure 5A by replacing the enhancer conservation count annotation in the joint model with the binary conserved enhancer annotation, and confirmed that this did not significantly change our results (Table S21). Second, we repeated the analysis of Table S21 by including 500bp flanking regions around each of the annotations from Figure 5A. We confirmed that this did not significantly change our results; the enrichment for conserved enhancer was slightly reduced, but remained highly significant (Table S22). Third, we repeated the analysis of Figure 5B by including human-specific promoters and promoter conservation count as additional annotations, in order to investigate whether this might lead to a significant *τ*^*^ for the promoter conservation count annotation (as in Table S15) due to the parabolic trend for promoters in Figure 3A. However, the *τ*^*^ for both annotations was non-significant (Table S23).

## Discussion

Our results helps elucidate which regulatory elements make the largest contributions to the genetic architecture of diseases and complex traits. We reached three main conclusions. First, disease heritability is concentrated in enhancers and promoters with ancient sequence age. Second, disease heritability is concentrated in enhancers and promoters with conserved function across species. Third, disease heritability is concentrated in promoters of ExAC LoF intolerant genes. These findings represent unique information about disease heritability conditional on all other available annotations, as quantified by large and highly significant *τ*^*^ values (up to 0.57 in Combined joint model; Figure 5B), substantially larger than the *τ*^*^ values that we reported for other annotations in our recent work ^7,26^. In addition to improving our biological understanding of disease architectures, our findings have immediate downstream applications to improve association power ^3,34,35^, fine-mapping ^2,36,37^, and genetic risk prediction ^38^–^40^.

Promoters are known to be functionally conserved more often than enhancers ^9^; we determined that conserved enhancers, although less common than conserved promoters, are particularly strongly enriched for disease heritability (Figure 5). In addition, previous work reported that human-specific DHS were significantly enriched for disease-and trait-associated variants, despite decreased constraint; we observed modest enrichment for human-specific promoters but no enrichment for human-specific enhancers (Figure 3A). The excess enrichments for enhancers and promoters with ancient sequence age raises the question of whether genomic regions with ancient sequence age are broadly important; however, ancient sequence age was not conditionally significant in our analyses (Table S7). Our finding of increased disease enrichment in promoters of ExAC LoF intolerant genes ^12^ (Figure 5A) is consistent with evidence from eQTL studies ^26^; however, our promoter of ExAC gene annotation remains uniquely informative conditional on the annotations from ref. ^26^ (Table S19). Our finding of increased disease enrichment in promoters of ancient genes (Table S16) is consistent with previous work showing that genes linked to human disease are more often ancient than recently evolved ^18^; however, we determined that the promoter of ancient genes annotation was not uniquely informative once the promoter of ExAC genes annotation was included in our model. All of our findings are consistent with the action of negative selection on genetic variants that impact disease ^7,41^–^44^.

We note several limitations of our work. First, we focused our analyses on common variants by using a 1000 Genomes LD reference panel, but future work could draw inferences about low-frequency variants using larger reference panels ^44^. Second, our main analyses were restricted to enhancers and promoters identified in liver tissue ^9^. Results involving sequence age were similar for other enhancer and promoter annotations (Table S8). However, efforts to generalize our results for conserved function are limited by the availability of enhancer and promoter annotations across species in other tissues; one possible solution would be to predict regulatory function across species in other tissues ^45^–^52^. Third, inferences about components of heritability can potentially be biased by failure to account for LD-dependent architectures ^7,53^–^55^. All of our analyses used the baseline-LD model, which includes 6 LD-related annotations ^7^. The baseline-LD model is supported by formal model comparisons using likelihood and polygenic prediction methods, as well as analyses using a combined model incorporating alternative approaches ^56,57^; however, there can be no guarantee that the baseline-LD model perfectly captures LD-dependent architectures. Despite these limitations, our results are highly informative for the genetic architecture of diseases and complex traits.

## Supplemental Data

Supplemental Data includes one figure and 23 tables.

## Acknowledgments

We are grateful to P. Flicek, P. Provero, D. Marnetto, H. Finucane, J. Stamatoyannopoulos, C. Breeze, and J. Vierstra for helpful discussions. This research was funded by NIH grants U01 HG009379, R01 MH101244, R01 MH107649 and 5T32CA009337-32. This research was conducted using the UK Biobank Resource under application 16549.

## Web Resources

All summary statistics from UK Biobank that we analyzed are publicly available at https://data.broadinstitute.org/alkesgroup/UKBB/

List of human genes that have a mouse ortholog in hg38 build: http://useast.ensembl.org/biomart/martview/ ^22^.

List of TSS of human genes in hg19 build: https://sourceforge.net/projects/seqminer/files/Reference%20coordinate/^23^.

Main annotations described in this manuscript are publicly available at https://data.broadinstitute.org/alkesgroup/LDSCORE/hujoel/.

